# Combined Linkage Disequilibrium and Linkage Analysis (cLDLA): implementation of a powerful approach to identify the genetic basis of complex traits in a bioinformatics workflow

**DOI:** 10.1101/2024.10.18.619071

**Authors:** M. Upadhyay, A. Panchal, I. Medugorac

## Abstract

Identifying the relationship between the polymorphism segregating in a population and phenotypic differences of a trait observed between the individuals of a population is of major biological interest and represents the basis of forward genetics. Much of the traits of interest are influenced by several polymorphic genes and environmental conditions. Often the loci associated with such measurable traits are referred to as Quantitative trait loci. These loci are identified using several statistical approaches. One of them is combined linkage disequilibrium and linkage analysis (cLDLA). This approach, first proposed by Meuwissen and colleagues in 2002, is shown to be robust against population stratification/family structure and requires a relatively lower sample size compared to a genome-wide association study design. Previously, we have successfully used this approach in mapping several important traits in livestock such as identifying the genetic basis of polled condition in cattle and tail length in sheep. A cLDLA requires several complex computation processing and intermediary file conversion steps; for some of these steps no open-source tools are available. Therefore, running this analysis, manually, can prove challenging, tedious, or error-prone. We present, cldla, a bioinformatics workflow implemented in nextflow which takes the vcf file and phenotype file as inputs and implements all the downstream processing required for cLDLA. Additionally, it also has a separate workflow to estimate SNP-based heritability and features for interactive visualization of the results. The workflow is freely available at: https://github.com/Popgen48/cldla.

## 1. Introduction

One of the important goals of modern biology is to understand how particular genotypes, whether alone or by interacting with other genotypes, give rise to a specific set of phenotypes. Usually, phenotypes of interest such as variation in height, weight, or disease conditions in humans, or milk production, fertility traits, and tail length in livestock, are the results of complex interactions between a large number of genes and multiple environmental conditions [1-3]. The chromosomal regions containing genes associated with such measurable traits are referred to as Quantitative trait loci (QTL) [4, 5]. Broadly, the most frequent methods used to identify QTL can be categorized into these two classes: (1) approaches based on genetic linkage mapping (also known as linkage analysis, LA), and (2) approaches based on linkage disequilibrium (LD) concepts like genome-wide association study (GWAS) [6, 7].

In human genetics, the classical approach was the application of parametric LA in families with clear disease inheritance e.g. [8, 9]. In agriculture and animal models the methods based on LA involve crossing the two divergent inbred lines to produce the F1 generation; the F1 generation is crossed to each other to produce the F2 or backcrossed with one of the parental lines to produce the BC1 generation. Subsequently, the phenotypes and genotypes of the F2 or BC1 generations are scored and likelihood ratios, which are based on statistical techniques such as composite interval mapping, are assessed to identify the chromosomal regions associated with the phenotype of interest [6]. This LA-based method has proven to be robust in identifying QTLs where a few loci have a large effect on the phenotype of interest. This method, however, is always not feasible as it can be expensive and lengthy. Moreover, the results obtained from such a design are not always applicable to general populations because of factors such as complex genetic architecture and phenocopies [5, 6].

On the other hand, studies like [10] proposed that the non-random association of alleles at different loci could be used to analyze the entire genome for disease loci. Houwen and colleagues were amongst the first to [11] provide empirical evidence that such non-random association, i.e. LD, can be used to map a human autosomal recessive disease gene by genome-wide screening. The GWAS for complex traits is based on the concept of LD, where the marker alleles associated with the causative variant are traced in unrelated affected individuals, who often share the identical haplotype descended from a common ancestor ten of generations back. In GWAS, thousands of genomic markers (usually SNPs) are genotyped and the statistical association between each of the markers and a phenotype is investigated using methods such as chi-squared test and generalized linear regression [3] based statistical models. In contrast to the LA-based methods, GWAS is carried out in an outbred population that has a similar ancestral composition [3]. It is a powerful method to detect common alleles having moderate to small effects on the phenotype. Compared to LA-based methods, it can also detect QTL at much higher resolutions because it exploits the historical recombination events. Precisely for this reason, the linkage disequilibrium (LD) decays fast even between those markers that are present over short genetic distances [5, 6]. Therefore, a large number of SNPs need to be genotyped to capture the association between QTL and genotyped markers. Further, GWAS suffers from limitations such as the confounding effect of population structure, population demography, family structure, cryptic relatedness, and requirement of large sample size [5, 12, 13].

By combining the information of LA and LD mapping into a single mapping design, in 2002, Meuwissen and colleagues proposed another approach to detect QTL [14]. Unlike the statistical association between each SNP and a phenotype as explored in GWAS, here, QTL is detected by estimating its variance components at each user-defined genomic window and contrasting it against the variance component associated with the background genes (no QTL) [14, 15]. This approach uses the phased genotypes (as opposed to a single marker in GWAS) which are substituted as random QTL effect in the flexible linear mixed model. Further, this mixed model can consider any pedigree structure by estimating genetic component variance based on identity-by-descent (IBD) covariances between individuals at each putative QTL location. The variance estimation is carried out via the restricted maximum likelihood approach (REML) [16] as implemented in *ASREML* [17] or using average information REML (AIREML) [18] as implemented in *blupf90+* [19]. This approach, combined linkage disequilibrium and linkage analysis (cLDLA), accounts for all recombination events since the foundation of a population, thereby, allowing researchers to estimate the genetic relationships among the individuals more accurately. The combination of LA and LD offers robustness against false positives because the signals must satisfy the assumption of both approaches. Although not widely used, the studies have indicated that cLDLA can detect QTL even when the sample size is relatively low or/and the phenocopies are present in the data [1, 20, 21]. For example, Lagler and colleagues [1] found that in their design of 362 animals which were genotyped for about 45,000 SNPs to find the QTL related to tail length in the Merinolandschaf breed, traditional GWAS didn’t show any strong signal with the genotyped SNPs. However, cLDLA identified the QTL in the upstream of the *HOXB13* gene. On the other hand, Gehrke and colleagues [22] showed that cLDLA is able to map autosomal recessive disorder in designs where the number of cases is severely limited, e.g. in a single half-sibling family with 15 affected and 12 unaffected offspring.

Despite these advantages, the cLDLA method is not used frequently because it is more complex and computationally intensive than traditional GWAS [23]. Moreover, to our knowledge, no open-source tools are available to perform the intermediate steps (described in the method section) required for the analysis. Further, compared to GWAS, cLDLA workflow takes longer to complete and there are very few specialized open-source software that are similar to ASREML to estimate the variance components. For these reasons, cLDLA is mostly used as a follow-up of GWAS to fine-map the target QTL or to remove the spurious signals once the putative regions encompassing QTLs are found [23, 24]. However, such follow-up requires successful initial mapping by GWAS, which in turn relies on large populations.

In the past, efforts were carried out to make the cLDLA accessible to the research community, for example, by incorporating it into a user-friendly web interface [5]. However, this web interface is not active anymore and, to our knowledge, currently no such alternative tools are available. To this end, here we present *cldla*, a workflow implemented in the Nextflow workflow management system to detect QTL using the cLDLA method [25].

## 2. Methods and implementation

### 2.1 Phasing of genotypes

The first step in the cLDLA is to phase the genotypes; for this purpose, Beagle (v 5.4) [26] and SHAPEIT5 (v 5.1.1) [27] are implemented in the workflow. Usually, it is recommended to filter out the non-informative markers before using the phased data for the estimation of IBD probability. Thus, the workflow removes the SNPs with the minor allele frequency (MAF) less than 0.025 by default. However, the user has the option to change this default or provide a vcf file that is already phased in a dataset that is much larger than the mapping population of interest. For the downstream processing, the workflow will only select the samples that are provided in the phenotype file. At a minimum, the phenotype file supplied to the workflow should contain the sample index, sample IDs, and the record of phenotype of interest for each of these samples.

### 2.2 Construction of mixed linear models (MLMs)

The essential components of the cLDLA approach are the mixed linear models (MLMs). It is assumed that a phenotypic trait of interest is controlled by a linear combination of fixed effects, putative QTL effect, and additive polygenic effect [15]. The additive polygenic effect and putative QTL are assumed to be random. Usually, such MLM is defined as:

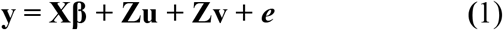

where **y** is a vector of phenotypes, **β** is a vector fixed effect. The vector **u** is of random additive polygenic effect with **u**∼N(0, **A**σ^2^_u_), where **A** is the additive genome-wide relationship matrix and σ^2^_u_ is the additive polygenic variance component. The vector **v** is of additive QTL effect with **v**∼N(0, **G**σ^2^ >_v_), where **G** is the covariance matrix of the additive effect of QTL and σ^2^ >_v_ is the additive variance of QTL. The vector **e** is of residual error with e∼N(0, **I**σ^2^_e_) where **I** is an identity matrix and σ^2^_e_ is the residual variance. The vectors **e, u**, and **v** are assumed to be uncorrelated.

Another MLM is constructed without assuming the QTL effect. It can be defined as:

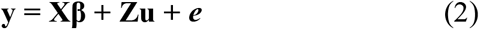

with **u**∼N(0, **A**σ^2^_u_) and e∼N(0, **I**σ^2^_e_).

At each QTL position (q), the parameters are estimated for (1) using a specialized tool such as ASREML or BLUPF90+. The parameters for model (2) are estimated once per chromosome or once per permutation (see below). Subsequently, the log-likelihood value (L1_q_) obtained for each QTL position is compared against the log-likelihood value (L0) obtained for (2) using the likelihood ratio test statistic (LRT):

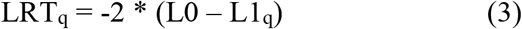

This general procedure of constructing MLMs and calculating LRT has already been described previously [14, 15]. The support for ASREML and BLUPF90+ tools are implemented in the workflow over other tools because of their ability to handle large user-defined (co)variance matrices or the inverse of such matrices.

To empirically estimate the genome-wide significance threshold, the distribution of LRT values under the null hypothesis is determined by a permutation test which is also implemented in the workflow. Briefly, for each chromosome, a user-defined number (default is 100) of windows are randomly selected; for each of these windows, the phenotypes are randomized, and the parameters of MLMs (1) and (2) are estimated. The highest LRT value obtained from this step is selected as the significance threshold to identify the putative QTL.

Alternatively, the users also can choose to obtain the P-values based on LRT values by using a χ2 distribution with one degree of freedom. The P-values threshold can be determined by a Bonferroni correction of the multiple testing (0.05/the number of non-overlapping windows) [28]. Here, the total number of non-overlapping windows are calculated as the number of SNPs divided by the window size for each chromosome separately.

### 2.3 Estimation of genetic relationship across the genome

In the equations (1) and (2), the polygenic variance (**A**) is estimated using the genomic relationship matrix (GRM). The inclusion of GRM corrects for the effect of population stratification and familial structure, thereby, reducing the spurious QTL signals. In cases, where the true candidate region is also included in the QTL as one of the effects of an MLM, the result may lead to a loss of power. Therefore, it is recommended to follow the “leave-one-chromosome-out” (LOCO) strategy [29]. In the context of cLDLA, the GRM can be estimated separately for each chromosome by excluding the genotyping data of the chromosome being assessed to identify QTL. By following this approach, the confounding covariance of the background genes on the putative QTL can be reduced. In the workflow, the “G.matrix” function of the *SNPReady* package is wrapped inside an R script [30]. This script estimates GRM using the methods of [31] or [32].

### 2.4 Estimation of genetic relationship at putative QTL

In equation (1), the additive genetic covariance at the putative QTL (G) is estimated by calculating IBD probabilities between all individuals at specific genomic positions. Several methods have been proposed to estimate the IBD probabilities using the genotype information with or without the pedigree data [20, 33-35]. In the workflow, the haplotype-based method proposed by [33] is implemented. This method accounts for multi-locus LD between the surrounding markers and the putative QTL position. Briefly, the IBD probability between any two haplotypes is estimated based on the identity and length of the marker haplotypes surrounding the putative QTL position [14, 33]. The Fortran module provided in the [33] paper was integrated into the C program estimating haplotype IBD probability at putative QTL for all haplotype combinations of all pairs of individuals in the mapping design.

In the workflow (Fig. 1), first, the phased genotypes are divided in the window of “n” SNPs (default value: 40) with a step size of one SNP. The position of putative QTL in each window is assumed as the mid-point of the same window similar to [14]. The IBD between any two haplotypes is therefore estimated based on the length of the allele chains, which are identical in both directions from this mid-point. The module considered allele and recombination frequency, i.e. the genetic length and marker density of the common IBD segment around the putative QTL to determine the level of IBD probability between two haplotypes. Next, this haplotype IBD is converted into a diplotype relationship matrix (DRM) based on the additive genotype coefficient method described in [36]. Briefly, for every pair of individuals, we estimated four haplotype IBD probabilities at putative QTL: (i) maternal haplotype of individual 1 (m_1_) versus paternal haplotype of individual 2 (p_2_), (ii) m_1_ versus m_2_, (iii), p_1_ versus p_2_, and (iv) p_1_ versus m_2_. The sum of these four IBD probabilities divided by two present the relationship between two individuals at a specific position in the genome, i.e. diplotype relationship. Here, the term, diplotype refers to the unique combination of two haplotypes of each pair in a mapping population.

**Figure 1.**
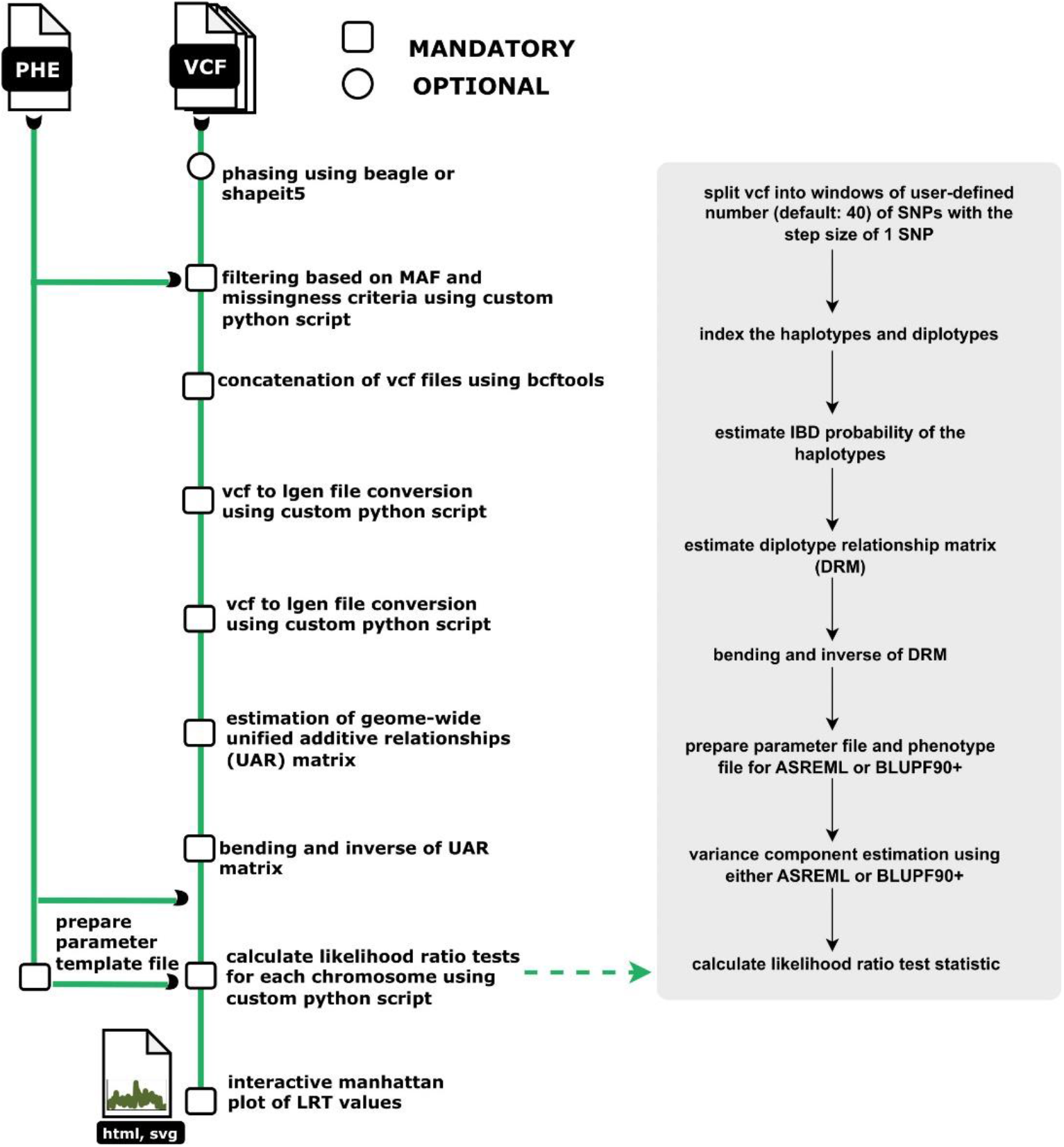
Overview of the cLDLA workflow. The workflow is mainly implemented using Nextflow workflow management system, Python, and Fortran programming languages. The steps highlighted along the green thick lines are implemented in Nextflow, while the steps highlighted in the thick gray box are implemented in python. Other significant steps such as bending and inverse of matrix as well as indexing the diplotypes use the Fortran programming language.

### 2.5 Bending and Inversion of GRM and DRM

In the workflow, the matrices — DRM of all the putative QTLs and GRM for each chromosome — are bent using the procedure as described in [37]. The Fortran codes provided by the same author were modified to reduce the computation time and supplied with the workflow. This bending procedure is required to ensure that the matrix is positive definite so that it can be inverted to find the solutions in MLMs. The inversion of matrices is carried out using the Fortran codes supplied with the package WOMBAT [38]. Finally, the inverted matrices are included in the estimation of parameters described in (1) and (2).

### 2.6 Estimation of heritability using genome-wide SNPs

The workflow also provides the option to estimate the phenotypic variance explained by genome-wide SNPs. To this end, genomic-relatedness-based restricted maximum-likelihood (GREML) as implemented in GCTA software version 1.92.3 is used [39]. Additionally, the permutation-test-based approach is also implemented in the workflow to assess the distribution of heritability estimates in the mapping designs under the null hypothesis [40].

Customized python script is implemented in the workflow to carry out interactive visualization of the results generated by the workflow.

## 3 Case study

### 3.1 QTL mapping of Tail length in Merinolandschaf breed

Lagler and colleagues [1] used the dataset of 362 phenotyped and genotyped Merinoland sheep to map the QTL affecting the tail length, a trait relevant to animal welfare in sheep breeding. In that study, the cLDLA was carried out by running several commands and scripts manually; further, as the input of one process was dependent on the output of another process, it was a tedious and time-consuming process. The workflow developed here replaces all these steps with a single command line. Therefore, to validate the workflow, we carried out the mapping again in the same design. In brief, the dataset consisted of 362 Merinoland sheep for which tail length, body weight (BW), height at withers (WH), sex, age and litter size were recorded. The animals were genotyped using the OvineSNP50 BeadChip from Illumina (Illumina, San Diego, USA) according to the manufacturer’s instructions. In the MLMs, BW, WH, gender, and age were used as fixed effects. The data was run through the workflow using the following command.

~~~
nextflow run popgen-cldla/ --input chrom_vcf_idx.csv –maf
0.025 --tool asreml -profile singularity --pheno_file TailMLS04.template.phe --outdir
TailMLS04_blupf90 --outprefix
TailMLS04_b -resume
~~~

To compare the results, the variance component estimation (VCE) was carried out using both, ASREML (version 4.2.1) and BLUPF90+ (version 2.51).

~~~
nextflow run popgen-cldla/ --input chrom_vcf_idx.csv –maf
0.025 --tool blupf90 -profile singularity --pheno_file
TailMLS04.template.phe --outdir TailMLS04_blupf90 --outprefix
TailMLS04_b -resume
~~~

The workflow generates the manhattan plot in terms of LRT values as well as in terms of P-values. The P-value threshold, as determined by the Bonferoni correction, is essentially the same for BLUPF90+ and ASREML approach. The numbers of SNPs satisfying this threshold, however, are different between BLUPF90+ and ASREML approach. Despite this, the potential candidate QTL on chromosome 11 as identified in validated in previous studies are detected by both these tools[1, 41]. The LRT values estimated from both tools yielded similar results (Figure 2). The significance threshold, however, as determined by the permutation tests are different. For the top 50 intervals on chromosome 11 having high LRT values (decided based on the BLUPF90+ threshold) the average difference in the LRT values between ASREML and BLUPF90+ was 0.23 (± 2.017) (Table S1). Note that the comparison was carried out after replacing the negative LRT values with zero in the outputs of ASREML and BLUPF90+. It should be noted that in the study of Lagler et al. (2022)[1], the interval with the highest LRT value was located at 37,111,462 bp on OAR11, which is as well among the intervals having the high LRT values as estimated by the workflow (Table S2). The highest LRT value as estimated by both the tools was at the interval 37,311,842 on chromosome 11. Interestingly, the causal mutations for the short-tail length in Merino sheep as proposed in the previous studies [1, 41] are closer to this interval (37,311,842 bp). The minor differences in the results could be due to the following factors: (i) Although we used the same genotypes as in Lagler et al., (2022), haplotyping with the Beagle program can lead to slightly different haplotypes. It is noteworthy that haplotyping is more accurate when performed in as large a population as possible, although only a small subset is used as a mapping population. After 2022, more sheep samples were genotyped with the OvineSNP50 BeadChip and used for improved haplotyping.

**Figure 2.**
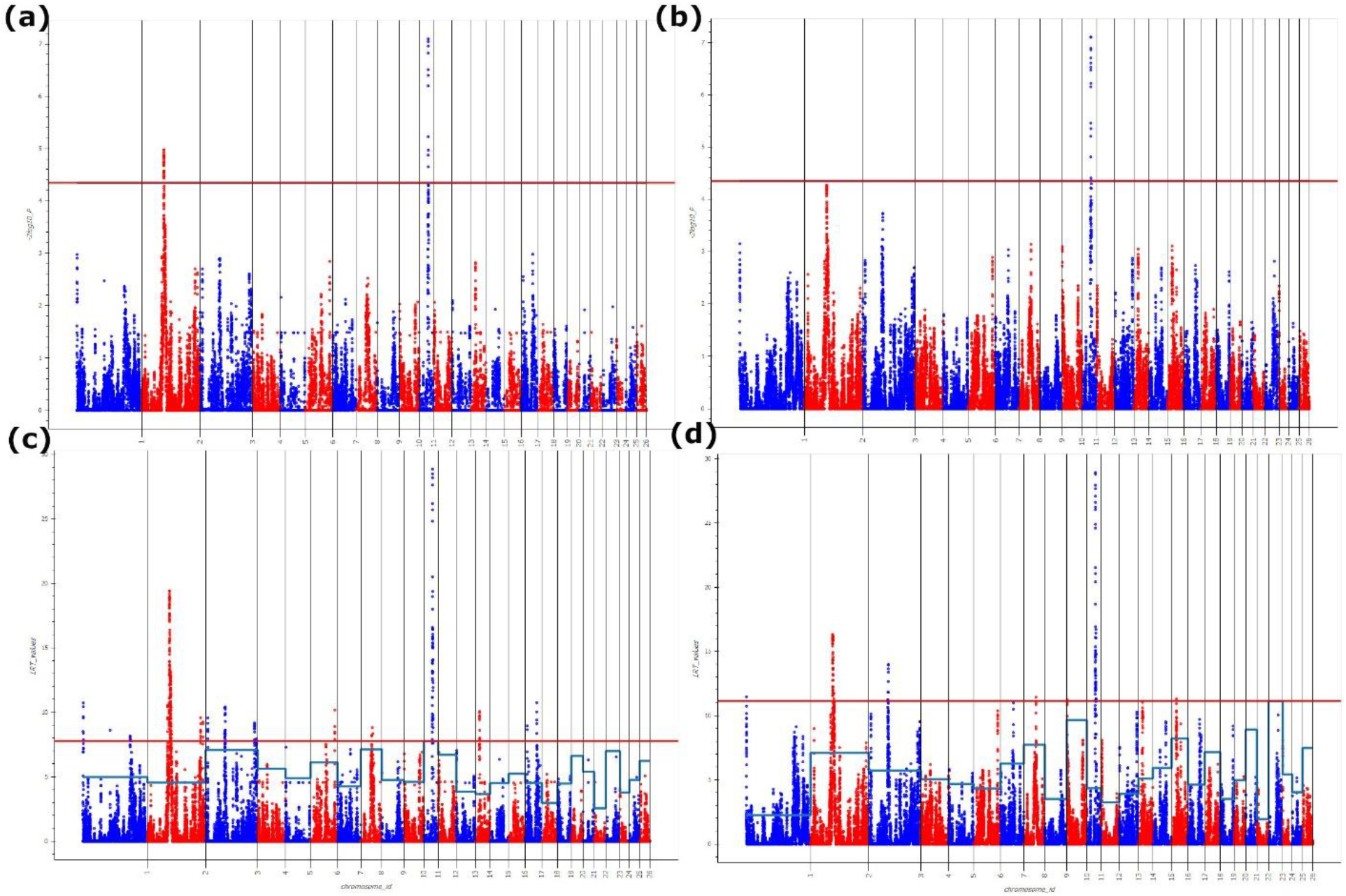
Manhattan plot visualization: -log10 of P-values calculated using (a) ASREML and (b) BLUPF90+, LRT values calculated using log-likelihood values of the MLMs estimated using (c) ASREML and (d) BLUPF90+. In case of a) and b), the red horizontal line represents the significant threshold for P-values after multiple correction testing, while in c) and d), it represents the highest LRT value observed across the chromosomes after a total of 2600 permutations (100 * 26). The blue zig-zag lines represent the highest LRT values observed for each chromosome after 100 permutations.

## 4. Discussion

In the past, cLDLA has been successfully used in our group to map the loci associated with various traits in livestock such as qualitative autosomal recessive and dominant traits in cattle as well as quantitative traits like tail length and ear length in sheep among others [1, 21, 40, 42-44]. Here, we packaged the various software developed in our group, in addition to the available open-source tools, using the container technology (docker and singularity). Further, we inter-connected these tools using the Nextflow workflow management system to implement cLDLA.

The current implementation of the cLDLA workflow has several limitations. For variance component estimation, the workflow supports ASREML (version 4.2.1) and BLUPF90+ (version 2.51). It should be noted that ASREML is licensed software, whereas BLUPF90+ is free only for research purposes. Additionally, for binary traits, the workflow currently supports variance component estimation using ASREML exclusively. Furthermore, no other random effects, apart from that of QTL itself, are supported in mixed linear models (MLMs). Despite these limitations, the workflow offers significant advantages, such as relatively small sample size requirements, robustness against population structure, and ease of analysis, making it a valuable tool for the research community engaged in mapping monogenic and polygenic traits in populations with a wide range of known and cryptic relationships.

